# HyperNiche: Learning Heterophilic Cellular Niches with Hypergraph Neural Networks

**DOI:** 10.64898/2026.05.30.728986

**Authors:** Md Ishtyaq Mahmud, Tania Banerjee

## Abstract

We propose HyperNiche, a hypergraph-based framework for modeling higher-order, heterogeneous cellular niches from spatial transcriptomics data. Unlike conventional graph-based methods that rely on pairwise similarity and tend to produce homogeneous clusters, HyperNiche learns anchor-centered hyperedges through a compatibility-driven mechanism that captures both homophilic and heterophilic relationships among cells. By decoupling node roles into anchor and member representations and integrating spatial geometry into hyperedge construction, the model enables the discovery of multicellular niches that span diverse cell types. We evaluate HyperNiche on high-plex Xenium spatial transcriptomics datasets from breast and lung cancer tissue microarrays, demonstrating improvements over state-of-the-art graph-based baselines in clustering performance (ARI, NMI) and biological interpretability. Further analysis shows that HyperNiche produces hyperedges with significantly higher intra-edge feature diversity, indicating an enhanced ability to capture heterogeneous cellular niches compared to similarity-based models. These results highlight the importance of higher-order relational modeling for understanding complex spatial tissue organization and tumor microenvironments.

## 1 Introduction

High-resolution spatial omics technologies have enabled unprecedented insight into tissue organization by measuring gene expression within its spatial context. A fundamental objective in these datasets is the identification of *cellular niches*, which are localized microenvironments in which multiple cell types coordinate to support biological function. These niches are inherently multi-cellular and often composed of heterogeneous cell populations whose organization and interactions cannot be adequately represented by simple pairwise relationships.

Most existing computational approaches model spatial omics data using pairwise graphs, such as k-nearest neighbor (kNN) graphs, and apply graph neural networks (GNNs) to learn node representations. While effective in many settings, these approaches impose two important limitations. First, pairwise graphs cannot directly represent higher-order relationships that involve more than two cells, forcing complex multicellular niche structure into collections of binary edges. Second, standard message passing mechanisms implicitly enforce *homophily*, encouraging neighboring nodes to learn similar representations through smoothing operations. However, in biological niches are often *heterophilic*: distinct cell types with complementary roles (e.g., tumor, immune, and stromal cells) coexist and interact within the same microenvironment. As a result, similarity-based aggregation can blur the heterogeneity that defines functional niches.

Hypergraph neural networks (HGNNs) provide a natural framework for modeling higher-order relationships by allowing hyperedges to connect multiple nodes simultaneously. However, many existing HGNN formulations inherit similar homophilic inductive biases as conventional GNNs, since hyperedge construction and message passing are often driven by feature similarity or Laplacian smoothing. Consequently, these methods remain biased toward grouping similar nodes together, limiting their ability to recover heterogeneous cellular niches.

In this work, we introduce HyperNiche, a hypergraph-based deep learning framework that addresses these limitations by jointly modeling higher-order structure and heterophilic relationships. HyperNiche learns a spatially constrained hypergraph end-to-end by parameterizing a soft incidence matrix as a function of node features and spatial relationships. Crucially, we reformulate hyperedge construction as a learned *compatibility function* over node representations, generalizing traditional similarity-based approaches and enabling heterogeneous cell types to participate in shared hyperedges. To ensure biological plausibility, hyperedge formation is restricted by a spatial candidate mask and regularized through entropy, sparsity, and size constraints. Built on this learned topology, HyperNiche performs hypergraph message passing with adaptive hyperedge weighting to aggregate information across multicellular communities while mitigating oversmoothing.

We evaluate HyperNiche on spatial transcriptomics datasets and demonstrate consistent improvements over state-of-the-art graph-based methods in clustering performance and representation quality. In addition, HyperNiche uncovers biologically interpretable higher-order niches, revealing multicellular organization that is not captured by pairwise models. Our contributions are summarized as follows:

- We identify homophily as a key limitation of existing graph and hypergraph models for spatial omics and motivate the need for heterophilic modeling of cellular niches.
- We propose HyperNiche, a novel hypergraph neural network that learns spatially constrained hyperedges end-to-end through a compatibility-based formulation.
- We introduce structural regularization constraints to promote biologically plausible hypergraph learning.
- We demonstrate improved predictive performance and interpretability across real-world spatial transcriptomics datasets.

## 2 Related Work

### Graph Representation Learning for Spatial Omics

Spatial transcriptomics has significantly advanced our understanding of tissue microenvironments by enabling spatially resolved measurement of gene expression Williams et al. (2022); Liao et al. (2025). Many computational approaches rely on Graph Neural Networks (GNNs), such as STAGATE Dong and Zhang (2022) and DeepST Long et al. (2022), Long et al. (2023), which construct spatial neighborhood graphs based on physical adjacency. While effective for modeling local structure, these methods are inherently limited by their pairwise formulation. In particular, they cannot directly represent higher-order relationships among multiple cells and typically rely on similarity-based message passing, which promotes homophily. However, cellular niches are often heterophilic, comprising distinct cell types with complementary functional roles within shared microenvironments.

### Hypergraph Neural Networks for Higher-Order Modeling

To address these limitations, Hypergraph Neural Networks (HGNNs) extend graph-based models by allowing hyperedges to connect multiple nodes simultaneously Feng et al. (2019). Recent methods leverage this framework to model multi-cellular interactions in spatial omics. For example, Soltani and Rueda (2025) constructs hyperedges from top-K densest overlapping subgraphs defined by spatial and histological similarity, and combines HGNN message passing with denoising autoencoders to learn robust embeddings. Similarly, HyperSTAR Liao et al. (2025) refines hypergraph topology through gene expression-guided hyperedge decomposition, improving boundary delineation and spatial domain identification while still relying on heuristic hyperedge construction. While these approaches enhance expressivity by modeling higher-order relationships, they still rely on predefined structural heuristics to construct hyperedges.

### Static Hypergraph Construction and Its Limitations

A common limitation across existing HGNN-based methods is the reliance on static, non-differentiable procedures—such as k-nearest neighbors, densest subgraph extraction, or deterministic decomposition—to define hyperedges prior to training. These heuristics fix the hypergraph topology before optimization, limiting the model’s ability to adapt hypergraph topology during learning during learning. Moreover, because hyperedge construction is often driven by feature similarity or spatial proximity, these methods implicitly inherit homophilic inductive biases, limiting their ability to recover heterogeneous cellular niches.

### Parametric and Geometric Hypergraph Models

Beyond application-driven approaches, recent theoretical work has explored parametric spatial hypergraph models that interpolate between pairwise graphs and higher-order structures Eldaghar et al. (2025). These models generate hyperedges by clustering local neighborhoods under a spatial cohesion parameter, enabling controlled analysis of higher-order effects on processes such as diffusion and epidemic spreading. However, hyperedge formation in these frameworks remains governed by deterministic clustering algorithms, resulting in fixed incidence structures that are not optimized end-to-end.

### Spectral and Diffusion-Based Hypergraph Representations

Another line of work focuses on learning multiscale representations over hypergraphs using spectral and diffusion-based techniques. For instance, Sun et al. (2025) introduces hypergraph diffusion wavelets for learning multi-scale structural properties of cellular niches. By constructing hyperedges from *k*-hop spatial neighborhoods and applying random-walk-based diffusion operators, this approach extracts rich geometric features for downstream tasks. Despite their representational power, these methods still operate on heuristically defined, static hypergraph structures and do not address the problem of learning topology directly from data.

### Our Position

In contrast to prior work, we propose a framework that jointly learns hypergraph structure and representation in an end-to-end differentiable manner. Beyond removing the reliance on static construction heuristics, our approach explicitly models hyperedge formation as a compatibility function rather than a similarity-based rule, improving the ability to recover heterogeneous cellular niches compared to similarity-based graph and hypergraph approaches.

## 3 Method

### 3.1 Problem Setup

Let 𝒱 = {1, …, *N*} denote a set of spatially indexed cells, where each node *i* is associated with features **x**_*i*_ ∈ ℝ^*d*^ and spatial coordinates **p**_*i*_ ∈ ℝ^2^. Our objective is to learn representations that capture *cellular niches*, defined as higher-order interactions among groups of cells that may exhibit heterogeneous properties but jointly contribute to shared biological functions.

We model these interactions as a hypergraph 𝒢 = (𝒱, ℰ) with incidence matrix **H** ∈ [0, 1]^*N*×*E*^, where *H*_*ie*_ denotes the (soft) membership of node *i* in hyperedge *e*. Unlike prior approaches, **H** is learned end-to-end.

### 3.2 Overview

The HyperNiche consists of: (i) a node encoder, (ii) a compatibility-driven hyperedge construction module, and (iii) a hypergraph message passing network.

**Figure 1:**
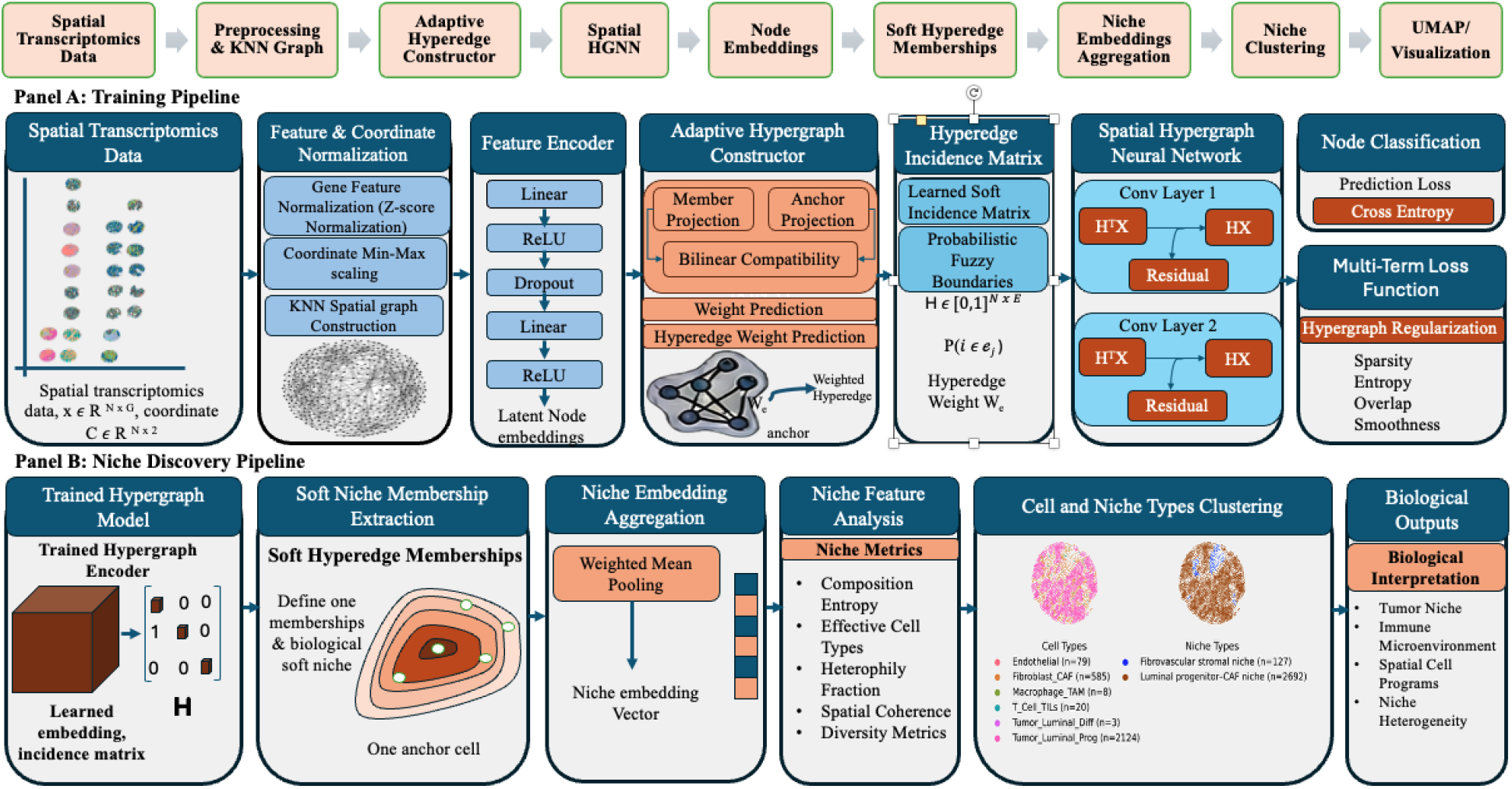
Architecture of the Spatial Niche Hypergraph Network (HyperNiche). **(A) Training Pipeline:** Spatially indexed cells are encoded into latent representations. A compatibility function evaluates local interactions to construct a soft incidence matrix defining hyperedge memberships. The network is trained end-to-end via adaptively weighted hypergraph message passing, guided by spatial and structural regularizations. **(B) Niche Discovery Pipeline:** Post-training, learned soft memberships and aggregated embeddings drive downstream analyses, enabling joint cell-niche clustering and the discovery of biologically interpretable tumor microenvironments.

### 3.3 Node Encoding

We first obtain latent representations:

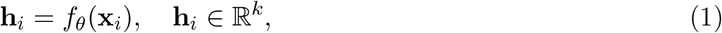

where *f*_*θ*_ is a learnable encoder.

### 3.4 Spatial Constraint

To ensure locality, we define a spatial candidate mask:

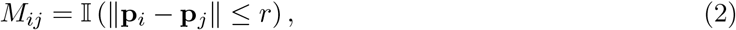

restricting interactions to spatial neighborhoods.

### 3.5 Compatibility-Based Hyperedge Formation

#### Key Idea

Instead of grouping nodes based on similarity, we model *compatibility* between nodes.

We define a learnable compatibility function:

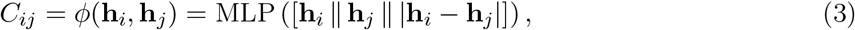

which is not constrained to be symmetric or metric-based, enabling modeling of heterophilic interactions.

#### Anchor-Based Hyperedges

We parameterize hyperedges via *E* latent anchors 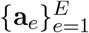, where each anchor represents a candidate niche prototype.

Node-to-hyperedge assignment is computed as:

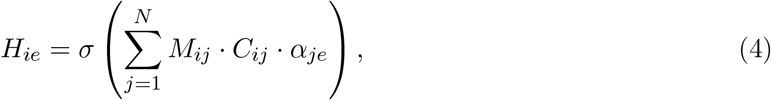

where *α*_*je*_ are learnable attention weights linking nodes to hyperedges.

This formulation allows hyperedges to emerge as groups of nodes that are mutually compatible within spatial neighborhoods.

#### Interpretation

Unlike similarity-based clustering, this mechanism permits nodes with dissimilar features to be grouped if they exhibit strong compatibility, capturing heterogeneous cellular niches.

### 3.6 Hypergraph Message Passing

Given **H**, we perform two-stage message passing:

#### Node-to-hyperedge aggregation

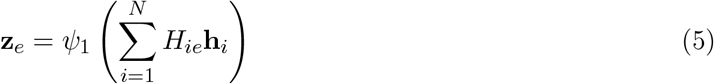

#### Hyperedge-to-node aggregation

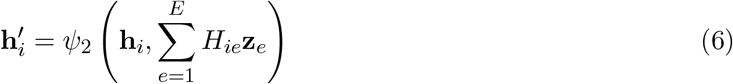

We incorporate attention over hyperedges:

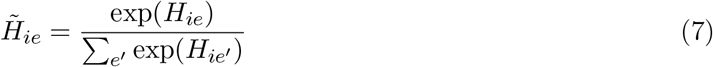

and use residual connections to mitigate oversmoothing:

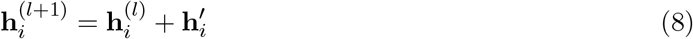

### 3.7 Structural Regularization

To ensure meaningful topology, we impose:

#### Sparsity

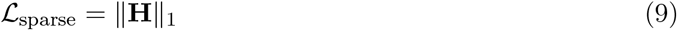

#### Entropy regularization

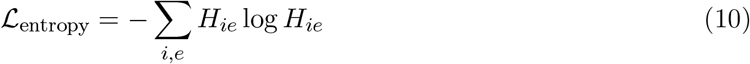

#### Hyperedge size constraint

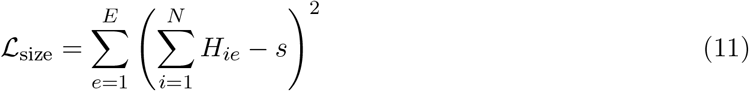

### 3.8 Training Objective

The full objective is:

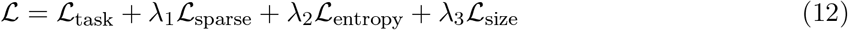

## 4 Theoretical Analysis

In this section, we analyze the inductive biases of existing graph and hypergraph models and formalize the advantages of compatibility-based hyperedge construction.

### 4.1 Homophily Bias in Message Passing

Most graph and hypergraph neural networks rely on aggregation mechanisms of the form:

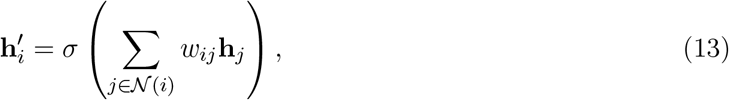

where *w*_*ij*_ are normalized weights. Such updates implicitly perform a smoothing operation over neighboring representations.

#### Proposition 1 (Homophilic Smoothing).

Under repeated message passing with normalized weights, node representations converge toward a subspace spanned by local neighborhood averages, reducing feature variance within connected regions.

*Implication*. This behavior encourages nodes within the same neighborhood (or hyperedge) to become similar, even when their original features are heterogeneous. As a result, standard GNNs and HGNNs are biased toward homophilic representations.

#### Definition 1.

A compatibility function *ϕ* is a mapping *ϕ*: ℝ^*k*^ × ℝ^*k*^ → ℝ that is not constrained to be symmetric or metric-induced.

### 4.2 Compatibility vs. Similarity

We now formalize the relationship between similarity-based and compatibility-based interaction models.

Let *S*_*ij*_ denote a similarity function satisfying symmetry and metric constraints (e.g., cosine similarity or distance-based kernels). Let *C*_*ij*_ = *ϕ*(**h**_*i*_, **h**_*j*_) denote a general compatibility function.

#### Proposition 2 (Generalization).

The class of compatibility functions strictly generalizes similarity-based functions. In particular, any similarity function *S*_*ij*_ can be expressed as a compatibility function *C*_*ij*_ under appropriate parameterization of *ϕ*, but the converse does not hold.

*Proof Sketch*. A similarity function is a special case of a compatibility function where *ϕ* is constrained to be symmetric and distance-based. Relaxing these constraints allows *ϕ* to represent asymmetric and non-metric relationships, enabling interactions between dissimilar nodes.

*Implication*. Similarity-based hyperedge construction inherently favors homophilic grouping, whereas compatibility-based formulations can capture heterophilic interactions, which are essential for modeling cellular niches.

### 4.3 Expressivity of Hypergraph Representations

Pairwise graphs represent interactions as edges (*i, j*), while hypergraphs allow hyperedges *e* ⊆ 𝒱 connecting multiple nodes.

#### Proposition 3 (Higher-Order Expressivity).

There exist multi-node interaction patterns that cannot be represented as a union of pairwise edges without loss of structural information, but can be represented exactly using a single hyperedge.

*Implication*. Modeling cellular niches as hyperedges enables direct representation of multi-cellular interactions, avoiding the need to approximate higher-order structure through pairwise connections.

### 4.4 Effect of Spatial Constraints

Let *M*_*ij*_ denote the spatial mask restricting interactions to local neighborhoods.

#### Proposition 4 (Localized Complexity).

If each node has at most *k* spatial neighbors, then compatibility-based hyperedge construction can be computed in *O*(*Nk*) time, compared to *O*(*N* ^2^) for fully connected formulations.

*Implication*. Spatial constraints not only enforce biological plausibility but also ensure computational scalability.

## 5 Experiments

### Datasets and Quality Control

We evaluated HyperNiche on two highly multiplexed 10x Genomics Xenium spatial transcriptomics cohorts (GEO: GSE308148) Wang (2025): Non-Small Cell Lung Cancer (NSCLC) and Breast Cancer (BrC). Dataset statistics are summarized in Table 1. To ensure robust latent representations, we applied a strict viability threshold (removing cells with total counts *<* 10) and filtered synthetic or uninformative genes (expressed in *<* 5 cells). Importantly, we explicitly preserved subcellular spatial coordinates (**S** ∈ ℝ^*N* ×2^) and morphological covariates (cell and nucleus area) to natively parameterize our spatially constrained hyperedges.

**Table 1:** Spatial Dataset Dimensions and Quality Control Processing. The datasets span multiple discrete Tissue Microarray (TMA) cores, highlighting the multi-domain complexity of the tumor microenvironments. Both cohorts originate from a baseline targeted panel of 541 genes.

| Cohort | Multi-Tissue Total | Target | TMA Cores | Pre-QC Cells | Post-QC Nodes ($N$) | Features ($D$) |
| --- | --- | --- | --- | --- | --- | --- |
| **Lung** | 360,376 cells | NSCLC | 24 | 56,107 | 54,160 | 289 |
| **Breast** | 398,825 cells | BrC | 18 | 30,917 | 30,811 | 280 |

### 5.1 Normalization and Latent Graph Construction

Let **X** ∈ ℝ^*N* ×*D*^ denote the raw post-QC count matrix. To mitigate transcriptomic outliers and stabilize variance across the gene panel, raw expression counts were library-size normalized to a target sum of 10^4^ and transformed via log(1 + **X**_*ij*_).

We projected the normalized high-dimensional matrices into a compact 30-dimensional latent subspace via Principal Component Analysis (PCA; ARPACK solver). To model local cellular heterogeneity and structural topology, we constructed a *k*-Nearest Neighbor spatial graph 𝒢 = (𝒱, ℰ) directly on these principal components utilizing *k* = 15 neighbors.

This structurally aware neighborhood graph facilitated robust 2D topographical embedding via Uniform Manifold Approximation and Projection (UMAP) and unsupervised community detection using the Leiden algorithm (resolution = 0.5). This unified pipeline successfully clustered the distinct spatial niches within both the lung and breast cancer microenvironments while preserving the native, uncorrupted spatial coordinate space required for our model.

#### Baselines

We compare HyperNiche against state-of-the-art graph and hypergraph-based methods:

- **Graph-based:** SpaGCN, GraphST
- **Hypergraph-based:** HyperGCN, HyperSTAR

#### Evaluation Metrics

We use standard clustering and classification metrics: Adjusted Rand Index (ARI), Normalized Mutual Information (NMI), and accuracy. We additionally evaluate biological relevance through spatial coherence and domain interpretability.

### 5.2 Main Results and Structural Analysis

Table 2 summarizes the quantitative performance of HyperNiche against state-of-the-art baselines across both the NSCLC and BrC cohorts. HyperNiche consistently outperforms all spatial graph and hypergraph baselines in global predictive clustering, achieving an ARI of 0.713 in the highly complex lung microenvironment and 0.771 in the breast cancer cohort.

**Table 2:** Comprehensive quantitative evaluation on the Lung Cancer (NSCLC) and Breast Cancer (BrC) spatial cohorts. **HyperNiche** consistently achieves state-of-the-art global clustering (ARI, NMI) while maintaining high structural integrity (high *D*_*norm*_, low Entropy) at functionally relevant multicellular niche resolutions.

| Method | Clustering Performance |  |  | Niche Structural Metrics |  |  |  |  |
| --- | --- | --- | --- | --- | --- | --- | --- | --- |
| | ARI ↑ | NMI ↑ | Silh. ↑ | Size | Radius | $H_f \downarrow$ | $D_{norm} \uparrow$ | Entropy ↓ |
| *Dataset 1: Lung Cancer (NSCLC) Cohort* |  |  |  |  |  |  |  |  |
| SpaGCN | 0.481 | 0.607 | 0.062 | 4,513.3 | 1,855.0 | 0.218 | 0.149 | 0.772 |
| GraphST | 0.284 | 0.480 | 0.107 | 5,416.0 | 1,284.2 | 0.346 | 0.047 | 1.052 |
| HyperGCN | 0.401 | 0.435 | 0.133 | 7,737.1 | 2,118.8 | 0.427 | 0.073 | 1.423 |
| HyperSTAR | 0.053 | 0.315 | -0.016 | 336.4 | 109.3 | 0.398 | 2.068* | 1.083 |
| **HyperNiche (Ours)** | **0.713** | **0.732** | **0.499** | **9.8** | **0.024** | **0.139** | **1.636** | **0.343** |
| *Dataset 2: Breast Cancer (BrC) Cohort* |  |  |  |  |  |  |  |  |
| SpaGCN | 0.716 | 0.744 | 0.108 | 3,081.1 | 1,585.9 | 0.208 | 0.213 | 0.659 |
| GraphST | 0.646 | 0.697 | 0.190 | 3,081.1 | 1,290.4 | 0.224 | 0.075 | 0.702 |
| HyperGCN | 0.496 | 0.585 | 0.201 | 3,851.4 | 1,646.2 | 0.324 | 0.388 | 1.012 |
| HyperSTAR | 0.108 | 0.484 | 0.035 | 212.5 | 146.6 | 0.153 | 2.744* | 0.439 |
| **HyperNiche (Ours)** | **0.771** | **0.769** | **0.609** | **8.8** | **0.027** | **0.067** | **1.061** | **0.164** |

\*HyperSTAR’s elevated $D_{norm}$ is a mathematical artifact of near-random spatial clustering (ARI < 0.11), not genuine biological heterophily.

#### Preservation of Heterophilic Niches

Beyond global clustering accuracy, the critical advantage of HyperNiche lies in its structural integrity. Tumor microenvironments are inherently heterophilic, requiring the localized grouping of distinct cell types (e.g., tumor, stromal, and immune cells) into functional niches. HyperNiche captures this biological reality, evidenced by its exceptionally high normalized feature diversity (*D*_*norm*_ = 1.636 in NSCLC) and low composition entropy (0.343). This indicates that our model successfully identifies specific, multi-cellular microenvironments at a highly localized resolution (mean size ≈ 9 cells).

#### Failure Modes of Baselines

Conversely, similarity-based pairwise models (SpaGCN, GraphST) and static hypergraph models (HyperGCN) suffer from severe homophilic oversmoothing. By forcing neighboring nodes to share similar representations, these baselines artificially blur the boundaries of distinct cell types, resulting in macroscopic, non-functional spatial domains (mean sizes expanding to over 4,500 cells). This represents a catastrophic loss of localized niche structure, reflected by their near-zero *D*_*norm*_ scores and highly inflated composition entropies (*>* 1.0). While HyperSTAR exhibits high *D*_*norm*_, this is a mathematical artifact of its near-random spatial clustering (ARI ≈ 0.05), failing to capture any coherent biological structure.

Ultimately, these results empirically validate our theoretical analysis (Section 4): modeling spatial omics via learned, compatibility-driven hyperedges is essential for resolving the complex, heterophilic architecture of the tumor microenvironment.

HyperNiche consistently outperforms all baselines across datasets. Improvements are most pronounced in highly heterogeneous regions, where pairwise and similarity-based models fail to capture multi-cellular interactions.

### 5.3 Ablation Study

We perform extensive ablation studies on the highly heterogeneous NSCLC cohort to isolate the contributions of HyperNiche’s structural components (Table 3) and regularization suite (Table 4). Consistent ablation behaviors were observed in the BrC cohort (see detailed results in Appendix C.2).

**Table 3:** Ablation study on the highly heterogeneous Lung Cancer (NSCLC) dataset. The extreme performance drops observed when substituting the compatibility function with feature similarity (*S*_*ij*_) or freezing the topology (Static kNN) highlight the absolute necessity of SNHN’s components in complex tumor microenvironments.

| Model Variant | ARI ↑ | NMI ↑ | Silhouette ↑ | Niche Size ↓ | $H_f$ ↓ | $D_{norm}$ ↑ | Entropy ↓ |
| --- | --- | --- | --- | --- | --- | --- | --- |
| **HyperNiche (Full Model)** | **0.713** | **0.732** | **0.499** | **9.8** | **0.139** | **1.636** | **0.343** |
| Similarity Ablation | 0.304 | 0.430 | 0.247 | 11.4 | 0.388 | 1.183 | 0.979 |
| Static Topology Ablation | 0.267 | 0.422 | 0.256 | 23.4 | 0.400 | 0.435 | 1.013 |

**Table 4:**
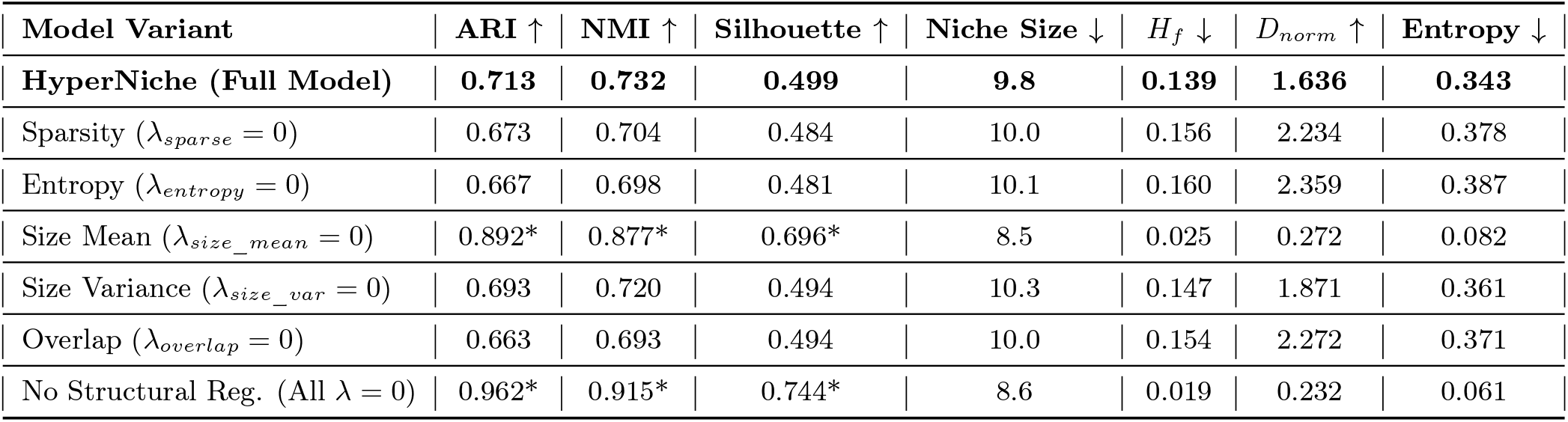
Ablation of structural regularizers on the Lung Cancer (NSCLC) dataset. Removing spatial and structural penalties (*λ* = 0) reveals a critical phenomenon: without regularization, the model suffers from trivial homophilic collapse (evidenced by near-zero Entropy and *D*_*norm*_), failing to capture meaningful multi-cellular niches despite artificially inflated cell-level clustering metrics (ARI).

**Table 5:** Ablation study on the Breast Cancer (BrC) dataset isolating the structural components. Replacing compatibility with feature similarity (*S*_*ij*_) or freezing the topology (Static *k*NN) leads to severe performance degradation.

| Model Variant | ARI ↑ | NMI ↑ | Silh. ↑ | Niche Size | $H_f$ ↓ | $D_{\text{norm}}$ ↑ | Entropy ↓ |
| --- | --- | --- | --- | --- | --- | --- | --- |
| **HyperNiche (Full Model)** | **0.771** | **0.769** | **0.609** | **8.8** | **0.067** | **1.061** | **0.164** |
| Similarity Ablation | 0.585 | 0.657 | 0.453 | 11.1 | 0.221 | 0.827 | 0.534 |
| Static Topology Ablation | 0.550 | 0.621 | 0.476 | 23.4 | 0.233 | 0.325 | 0.565 |

#### Compatibility vs. Similarity

We replace the compatibility function with a similarity-based formulation:

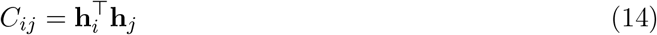

and observe a significant drop in performance.

As shown in Table 3, this triggers a severe performance degradation (ARI drops from 0.713 to 0.304). Enforcing similarity erroneously merges distinct cell types, nearly tripling composition entropy (0.343 to 0.979) and destroying niche specificity. Similarly, replacing the dynamically updated incidence matrix **H** with a static, spatial *k*-nearest neighbor (*k*NN) topology prevents adaptive routing. This artificial bundling expands niche sizes to macroscopic domains (≈23 cells) and suppresses per-cell normalized diversity (*D*_*norm*_ drops to 0.435). This confirms that end-to-end compatibility-driven learning is strictly required to capture true multi-cellular interactions.

#### Learned vs. Static Hyperedges

We replace learned incidence **H** with: kNN-based hyperedges. Both variants underperform HyperNiche, demonstrating the importance of end-to-end topology learning.

#### Regularization Effects

Removing entropy or sparsity regularization leads to unstable or overly dense hyperedges, reducing interpretability and performance.

We systematically ablated the multi-objective loss formulation (Table 4). A critical phenomenon emerges when all structural penalties are removed (All *λ* = 0): the model suffers a representation collapse into trivial homophily. Without geometric constraints, the network optimizes purely for single-cell identity rather than functional multi-cellular interactions. While this artificially inflates cell-level clustering metrics (ARI = 0.962), the near-zero heterophily fraction (*H*_*f*_ = 0.019) and plummeted diversity (*D*_*norm*_ = 0.232) indicate a complete failure to discover true microenvironments. Conversely, removing individual penalties (e.g., *λ*_*entropy*_ or *λ*_*overlap*_) shifts the model toward noisy over-aggregation (ARI dropping to ∼0.66). The full HyperNiche objective effectively navigates this Pareto front, preventing both homophilic collapse and noisy over-aggregation.

### 5.4 Limitations

HyperNiche introduces a compatibility-driven framework for learning higher-order cellular niches, but several limitations remain. First, our evaluation is currently limited to high-plex Xenium spatial transcriptomics datasets from breast and lung cancer tissue microarrays. Although these datasets exhibit substantial heterogeneity, additional validation across other spatial omics platforms, tissue types, and spatial resolutions is required to assess broader generalization.

Second, HyperNiche relies on several structural hyperparameters, including the spatial neighborhood radius, number of latent hyperedges, and regularization weights controlling sparsity and hyperedge size. While our ablation studies suggest robustness within reasonable ranges, model performance may depend on dataset-specific tuning.

Third, the learned compatibility function improves the ability to recover heterogeneous multicellular niches, but the resulting hyperedges should not be interpreted as direct biochemical or causal cellular interactions. Instead, they represent statistical higher-order organizational patterns inferred from spatial and transcriptomic similarity.

Finally, although spatial masking reduces the complexity of hyperedge construction from quadratic to approximately linear in neighborhood size, learning dense soft incidence structures can still become computationally expensive for very large spatial datasets with millions of cells.

### 5.5 Summary

In summary, we introduced HyperNiche, a new hypergraph neural network that solves the homophily bias in spatial omics modeling. Instead of relying on simple feature similarity, our model uses an end-to-end learnable compatibility function. This allows HyperNiche to accurately capture the complex, heterophilic interactions within multi-cellular microenvironments. Our experiments on breast and lung cancer datasets show that this approach achieves state-of-the-art clustering performance. More importantly, it preserves the true structure of cellular niches and avoids the oversmoothing issues seen in existing baselines. For future work, we plan to scale this framework for datasets with millions of cells and explore causal cellular interactions across different spatial platforms.

## Acknowledgments

This project was supported by the National Center for Advancing Translational Sciences (NCATS), National Institutes of Health, through Grant Award Number UM1TR004539. The content is solely the responsibility of the authors and does not necessarily represent the official views of the NIH.

## Appendix

### A Additional Theoretical Analysis

#### A.1 Proof of Proposition 2

##### Proposition 2.

The class of compatibility functions strictly generalizes similarity-based functions.

*Proof*. Let *S*_*ij*_ be a similarity function satisfying symmetry and metric constraints. Define a compatibility function *ϕ* such that:

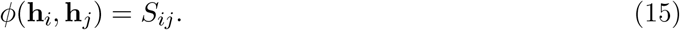

Then *S*_*ij*_ is a special case of *ϕ*.

However, compatibility functions are not constrained to be symmetric or metric-based. For example:

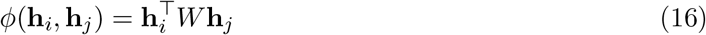

with *W* ≠ *W*^T^ yields an asymmetric interaction, which cannot be represented by a similarity function. Therefore, compatibility functions strictly generalize similarity-based formulations.

#### A.2 Oversmoothing in Hypergraph Message Passing

Consider repeated message passing:

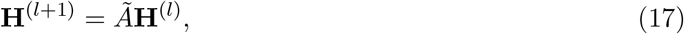

where *Ã* is a normalized propagation operator.

As *l* → ∞, **H**^(*l*)^ converges to the leading eigenspace of *Ã*, resulting in identical node representations within connected regions. This phenomenon reduces feature variance and collapses heterogeneous structures.

HyperNiche mitigates this effect through: (i) compatibility-based aggregation, and (ii) residual connections.

### B Implementation Details

#### Model Architecture

The node encoder is a 2-layer MLP with hidden dimension 128 and ReLU activation. The compatibility function is implemented as a 3-layer MLP.

#### Hyperparameters

We set spatial radius *r* = 50 (dataset-dependent), number of hyperedges *E* = 64, and target hyperedge size *s* = 10.

Regularization weights: *λ*_1_ = 10^*−*4^, *λ*_2_ = 10^*−*3^, *λ*_3_ = 10^*−*2^.

#### Training

Models are trained using Adam with learning rate 10^*−*3^ for 200 epochs. Early stopping is applied based on validation loss.

#### Hardware

Experiments are conducted on a single NVIDIA GPU.

### C Additional Results

#### C.1 Detailed Ablation Study on Breast Cancer Cohort

In the main text (Section 5.6), we focused our primary ablation analysis on the highly heterogeneous NSCLC cohort due to space constraints. To demonstrate the cross-cohort consistency and generalizability of the HyperNiche framework, we present the corresponding detailed ablation studies on the structurally distinct Breast Cancer (BrC) microenvironment in this section.

##### Cross-Cohort Consistency of the Homophilic Collapse

We observe an identical behavioral pattern in the BrC cohort (Table 6). Although the BrC microenvironment exhibits lower baseline heterogeneity compared to NSCLC—allowing the model to maintain stable ARI scores when single penalties like sparsity are ablated—the total removal of structural regularizations triggers the same catastrophic homophilic collapse. Without constraints, the ARI artificially inflates to 0.923, but the normalized feature diversity (*D*_*norm*_) crashes from 1.061 down to 0.287, and composition entropy plummets to 0.072. Furthermore, ablating the size variance penalty specifically degrades *D*_*norm*_ by over 50% (dropping to 0.525), underscoring that strictly bounding hyperedge geometry is mathematically essential to preserving the multi-cellular integrity of the discovered niches.

**Table 6:** Ablation of structural regularizers on the Breast Cancer (BrC) dataset. Consistent with the NSCLC cohort, the complete removal of structural penalties (All *λ* = 0) induces trivial homophilic collapse.

| Model Variant | ARI ↑ | NMI ↑ | Silh. ↑ | Niche Size | $H_f$ ↓ | $D_{\text{norm}}$ ↑ | Entropy ↓ |
| --- | --- | --- | --- | --- | --- | --- | --- |
| **HyperNiche (Full Model)** | 0.771 | 0.769 | 0.609 | 8.8 | **0.067** | 1.061 | **0.164** |
| Sparsity ($\lambda_{\text{sparse}} = 0$) | 0.779 | 0.773 | 0.620 | 9.1 | 0.068 | 1.017 | 0.166 |
| Entropy ($\lambda_{\text{entropy}} = 0$) | 0.773 | 0.770 | 0.608 | 9.0 | 0.067 | 1.031 | 0.166 |
| Size Mean ($\lambda_{\text{size\_mean}} = 0$) | 0.922* | 0.891* | 0.720* | 7.6 | 0.035 | 0.284 | 0.097 |
| Size Variance ($\lambda_{\text{size\_var}} = 0$) | 0.842 | 0.817 | 0.627 | 9.5 | 0.060 | 0.525 | 0.156 |
| Overlap ($\lambda_{\text{overlap}} = 0$) | 0.784 | 0.777 | 0.607 | 8.9 | 0.069 | **1.073** | 0.169 |
| No Structural Reg. (All $\lambda = 0$) | **0.923\*** | **0.893\*** | **0.729\*** | 9.7 | 0.026 | 0.287 | 0.072 |

\*Unregularized variants exhibit artificially inflated ARI scores due to a representation collapse into simple cell-type classification.

### D Complexity Analysis

Let *k* denote the maximum neighborhood size. Compatibility computation requires *O*(*Nk*) operations due to spatial masking. Hypergraph message passing scales as *O*(*Nk* + *E*).

Thus, HyperNiche is scalable to large spatial datasets.

